# Dietary environmental factors shape the immune defence against *Cryptosporidium* infection

**DOI:** 10.1101/2023.03.30.534739

**Authors:** Muralidhara Rao Maradana, N. Bishara Marzook, Oscar E. Diaz, Tapoka Mkandawire, Nicola Laura Diny, Ying Li, Anke Liebert, Kathleen Shah, Mauro Tolaini, Martin Kváč, Brigitta Stockinger, Adam Sateriale

## Abstract

*Cryptosporidium* is a leading cause of diarrheal-related deaths in children, especially in resource-poor settings. It also targets the immunocompromised, chronically infecting people living with HIV and primary immunodeficiencies. There is no vaccine or effective treatment. While it is known from human cases and animal models that CD4^+^ T-cells play a role in curbing *Cryptosporidium*, the role of CD8^+^ cells remains to be defined. Using a *Cryptosporidium tyzzeri* mouse model, we show that gut-resident CD8^+^ intraepithelial lymphocytes (IELs) confer resistance to parasite growth. CD8^+^ IELs express, and are dependent on, the ligand-dependent transcription factor aryl hydrocarbon receptor (AHR). AHR deficiency reduced CD8^+^ IELs, decreased their cytotoxicity, and worsened infection. Transfer of CD8^+^ IELs rescued severely immunodeficient mice from death following *Cryptosporidium* challenge. Finally, dietary supplementation of the AHR pro-ligand indole-3-carbinol to new-born mice promoted resistance to infection. Therefore, common dietary metabolites augment the host immune response to cryptosporidiosis, protecting against disease.

## INTRODUCTION

Diarrhoea contributes to nearly 11% of early childhood mortality worldwide (Liu et al., 2012). *Cryptosporidium* is an apicomplexan parasite that invades epithelial cells of the small intestine (Guerin and Striepen, 2020) and its infections are the second-leading cause of severe diarrhoeal events in young children in resource-poor regions (Kotloff et al., 2013). Recurrent infections are associated with malnutrition, leading to lasting effects such as growth stunting and impaired cognitive development (Kabir et al., 2021; Khalil et al., 2018; Molbak et al., 1997). *Cryptosporidium* is also an important opportunistic pathogen in immunocompromised individuals such as people living with HIV, transplant and chemotherapy recipients, and patients undergoing treatment for haemodialysis and cancer (Manabe et al., 1998; Sepahvand et al., 2022). Human cryptosporidiosis is usually caused by anthroponotic *Cryptosporidium hominis* or zoonotic *Cryptosporidium parvum* (Hossain et al., 2019). There is no vaccine against cryptosporidiosis. Nitazoxanide, the only FDA approved drug to treat cryptosporidiosis, has limited efficacy in immunocompetent individuals and is ineffective in HIV-AIDS patients and malnourished individuals (Amadi et al., 2002; Amadi et al., 2009). Although some progress has been made towards the development of new therapeutics (Manjunatha et al., 2017), this has been hindered by a lack of physiologically relevant model systems of cryptosporidiosis (Manjunatha et al., 2017; Marzook and Sateriale, 2020). We previously developed a mouse model of cryptosporidiosis with a natural mouse-infecting species, *Cryptosporidium tyzzeri,* which recapitulates the natural course of infection and intestinal pathology of human disease (Sateriale et al., 2019).

Increased susceptibility of patients with primary immunodeficiencies and experimental infections of human volunteers suggests an immune system-mediated protection from cryptosporidiosis (Cohn et al., 2022). Furthermore, most children living in endemic regions develop protective immunity to subsequent infections (Kabir et al., 2021; Kattula et al., 2017). Early studies on athymic mice revealed T-cells as important regulators of *C. parvum* infection (Heine et al., 1984). Increased infection burden in HIV-AIDS patients with low CD4+ T cell counts also highlights the importance of interferon-γ (IFNγ)-producing CD4^+^ T cells (Kaplan et al., 2009). Mice lacking T-cells (but not B cells) are unable to control a *C. tyzzeri* infection (Sateriale et al., 2019). Furthermore, mice without mature T and B cells, and those lacking IFNγ, cannot elicit vaccination-mediated protection during a secondary parasite infection (Sateriale et al., 2019). Mice harbouring a commensal strain of *C. tyzzeri* elicit both innate and adaptive immune responses, with early evidence hinting at elevated CD8^+^ T cells in infected mice (Russler-Germain et al., 2021). CD8^+^ T cells are highly enriched in the intestine and notably heterogeneous in phenotype and function. ‘Conventional’ mucosal T cells express TCRαβ together with CD4 or CD8αβ as TCR coreceptors and reside in the lamina propria. ‘Non-conventional’ mucosal T cells, expressing either TCRαβ or TCRγδ and typically also CD8αα homodimers, are prevalent in the mucosal epithelium (Hayday et al., 2001; Sujino et al., 2016). Thymic-derived natural intraepithelial lymphocytes (IELs) express CD8αα (TCRαβ^+^CD8αα, TCRαβ^+^CD4^+^CD8αα and TCRγδ^+^CD8αα), whereas peripherally induced IELs express CD8αβ (TCRαβ^+^CD8αβ) (Cheroutre et al., 2011; Hoytema van Konijnenburg and Mucida, 2017).

CD8^+^ IELs cells constantly scan epithelial cells for injury or infection and are considered primary responders to epithelial damage (Hoytema van Konijnenburg et al., 2017). CD8^+^ IELs are elevated in calves and mice infected with *C. parvum* (Chai et al., 1999; Pasquali et al., 1997). Nevertheless, factors that influence CD8^+^ IEL-mediated immunity and mechanisms by which CD8^+^ IELs confer resistance to *Cryptosporidium* are unknown. Most CD8^+^ T cells exhibit a tissue-resident memory phenotype (Sasson et al., 2020) and their maintenance depends on stimulation of the aryl hydrocarbon receptor (AHR) by endogenous ligands such as tryptophan-derived phytochemicals, microbial metabolites, and indole derivatives (Li et al., 2011) . AHR is a ligand-dependent transcription factor abundantly expressed in barrier tissues such as the gut, skin, and lungs. AHR is expressed by various cells in the gut including barrier epithelium, endothelium, and immune cells, and is hence a key factor in maintaining gut barrier integrity (Stockinger et al., 2021). AHR deficiency or depletion of AHR ligands increases susceptibility to bacterial infection in the colon (Stockinger et al., 2021) and also contributes to colon tumorigenesis (Metidji et al., 2018).

*Cryptosporidium* infects and replicates inside small intestinal epithelial cells, causing villus blunting and crypt hyperplasia and thereby significant gut damage (Sateriale et al., 2019). In human volunteers infected with *C. hominis* or *C. parvum,* increased fecal indole levels prior to infection correlated with decreased parasite burden (Chappell et al., 2016), indicating a protective function of indoles in cryptosporidiosis through an unknown mechanism. We wondered if these indoles were working via AHR-mediated gut protection.

To test this, we created a novel *C. tyzzeri* reporter strain expressing luminescent and fluorescent proteins utilizing an isolate of *C. tyzzeri* from the Czech Republic (*Ct*-CR2206, shortened here to *Ct*-CR) (Kvac et al., 2013). Here we show that in immunocompetent wild-type mice, *Ct-*CR altered epithelial differentiation and triggered an expansion of CD8^+^ IELs, and IELs conferred protection against infection when transferred to immunodeficient mice. Immune cell-specific deletion of AHR or the deprivation of AHR ligands in mice greatly depleted CD8^+^ IELs. Furthermore, dietary supplementation of AHR ligands to nursing mothers and their weaned pups provided prophylactic defence against infection. This highlights the opportunity for diet-based therapeutic interventions to treat this debilitating disease in humans and ruminant animals.

## RESULTS

### *Cryptosporidium tyzzeri* infects the ileum and affects epithelial cell differentiation

The *Cryptosporidium tyzzeri* strain Ct-CR2206 was originally isolated from a wild mouse in the Věrušičky municipality of the Czech Republic (Kvac et al., 2013). We sequenced the genome of *Ct-*CR2206 (shortened here to ‘*Ct-*CR’) using Illumina short read sequencing. Reads were mapped using the full *C. tyzzeri*-UGA55 genome as a reference (**Figure 1A**). In total, 13139 SNPs and 2983 insertion-deletion events were detected between the two strains (see **Supplementary Table S1** for a further breakdown). We genetically engineered *Ct*-CR to express the fluorescent mNeonGreen protein, along with a Nanoluciferase and Neomycin resistance cassette for easy parasite detection and transgenic selection, respectively (**Figures 1B and 1C**). Parasite replication over time could be tracked by assaying Nanoluciferase activity in mouse fecal pellets (**Figure 1D**).

**Figure 1:**
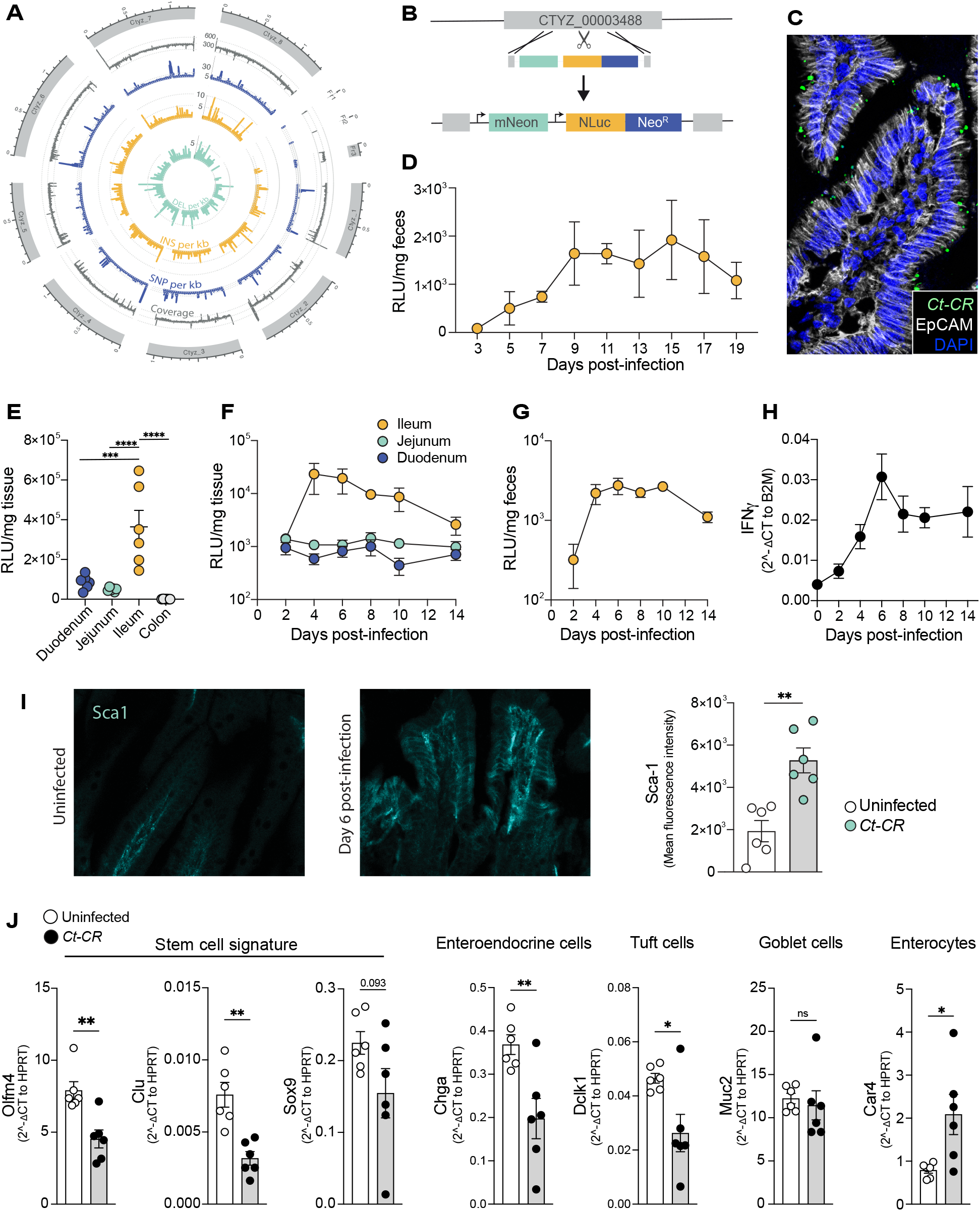
*Cryptosporidium tyzzeri* specifically infects the ileum and alters epithelial cell differentiation. (A) Chromosome-based Circos plot mapping the newly-sequenced *C. tyzzeri*- CR to the *C. tyzzeri*-UGA55 strain. Tracks from the outside-in are mean genome coverage per kb (grey), single-nucleotide polymorphism (SNP) density per kb (blue), insertions density per kb (orange), and deletions density per kb (green). (B) Schematic of the cloning strategy for the introduction of genes expressing mNeonGreen, Nanoluciferase (NLuc), and Neomycin phosphotransferase (Neo^R^) by Cas9-directed homology repair of the *C. tyzzeri*-CR Thymidine Kinase gene. (C) Immunofluorescence image showing mNeonGreen-expressing parasites in the intestinal villi from the ileal section of an immunocompromised mouse. (D) *C. tyzzeri* (*Ct*- CR) parasite burden in the feces measured by Nanoluciferase assay (E) Regional-specific *Ct-* CR parasite burden in infected wild type (WT) mice. (F) *Ct-*CR burden in the duodenum, jejunum and ileum. (G) Fecal levels of *Ct-*CR in infected WT mice. (H) qPCR of IFNγ in ileum tissue. (I) Sca-1 expressing epithelial cells in uninfected and *Ct-*CR infected WT mice. (J) qPCR of purified epithelial cells for marker genes of stemness (*Olfm4*, *Clu*, *Sox9*) and differentiated epithelial cells (*Chga*, *Dclk1*, *Muc2*, *Car4*). (F-H) N=4 per time point (E, I-J) Representative of at least 2 independent experiments. Each dot represents individual mice. RLU-Relative Luciferase Units. Error bars, mean + SEM. ns-not significant, *p < 0.05, **p < 0.01, ***p < 0.001, ****p < 0.0001 as calculated by t-test or one- way ANOVA with Tukey post-test.

*Ct*-CR infects the small intestine but not the colon **(Figure 1E)**. To determine which specific region in the small intestine harboured parasites, we infected C57BL/6 wild-type (WT) mice and measured luciferase levels from duodenal, jejunal, and ileal tissue every other day up to day 14 post-infection (DPI-14). This revealed the ileum to be the major site of *Ct-*CR expansion within the small intestine **(Figure 1F).** This infection location is similar to previously described *C. tyzzeri* strains and to what is seen during human cryptosporidiosis (Russler-Germain et al., 2021; Sateriale et al., 2019). Luciferase levels in fecal samples from the same mice reflected the parasite burden in the ileum **(Figure 1G)**. Therefore, we used readings from fecal samples as representative measures of the *Ct-*CR burden in the mouse ileum.

IFNγ is key to controlling *Cryptosporidium* growth, for both *C. parvum* (Griffiths et al., 1998; Gullicksrud et al., 2022; Robinson et al., 2001) and the previously isolated US *C. tyzzeri* strain, UGA55 (Sateriale et al., 2019). Consistently, infection with the *Ct-*CR strain led to a gradual increase *in Ifng* transcripts in the ileum **(Figure 1H)**, suggesting that the *Ct-*CR strain is pathogenic and induces inflammation in the infected area. Sca-1 (also known as Ly6a) is a marker of mouse intestinal epithelial injury, initially identified in the context of colitis (Yui et al., 2018) and staining of ileal sections showed a significant increase in Sca-1 expression in the villi of infected mice **(Figure 1I).** To study how *Cryptosporidium* changes the epithelial cell composition in the small intestine, we purified epithelial cells at the peak of infection (DPI-6) and compared their gene expression with that of uninfected controls. Epithelial cells from infected mice showed reduced expression of the stem cell signature (*Olfm4, Clu, Sox9*) and reduced expression of markers for enteroendocrine cells (*Chga*) and Tuft cells (*Dclk1*) **(Figure 1J).** However, no change was seen in expression of the goblet cell marker *Muc2* **(Figure 1J).** In contrast, there was an increase in the enterocyte marker (*Car4*) **(Figure 1J).** Therefore, the *Ct-*CR strain used in this study is pathogenic in nature and causes inflammation, epithelial cell injury and alters the cellular composition of the small intestinal epithelium.

### Haematopoietic cell specific aryl hydrocarbon receptor-deficient mice are susceptible to *C. tyzzeri* infection

AHR is a transcriptional regulator of genes involved in anti-microbial defence and intestinal epithelial differentiation. It has been shown to play a protective role during colonic bacterial infection and chemically-induced colon damage (Metidji et al., 2018; Shah et al., 2022; Stockinger et al., 2021). To determine whether AHR executes similar disease protective functions in the small intestine, we infected full body AHR knockout (AHRKO^Body^) mice and co-housed WT littermate controls with *Ct-*CR. AHRKO^Body^ mice displayed increased parasite burdens from the beginning and maintained high parasite loads throughout the course of infection (**Figure 2A** left panel), also assessed by area under the curve (AUC) (**Figure 2A** right panel). Having established the importance of AHR for *Cryptosporidium* infection control, we next wanted to know which cells were playing a role in this defence. Using AHR-reporter mice expressing AHR td-Tomato, we determined that the majority of epithelial cells express AHR (**Figure 2B)**, consistent with previous observations (Diny et al., 2022). Since *Cryptosporidium* only infects epithelial cells, we first challenged mice with an intestinal epithelial cell (IEC)- specific AHR knockout (Vil-Cre AHR^fl/fl^ termed AHRKO^Epithelium^) with *Ct*-CR to probe whether IEC-intrinsic AHR expression influenced parasite burden. Surprisingly, we found that epithelial AHR signalling was dispensable for control of parasite growth **(Figure 2C)**. AHR is also expressed by immune cells in the gut (**Figure 2D)**. We next asked whether AHR expression in immune cells is required to limit *Ct*-CR. To achieve this, we crossed hematopoietic cell-specific Vav-Cre mice with AHR^fl/fl^ mice to produce immune cell-specific AHR-deficient mice (AHRKO^Immune^). Consistently, mice with AHR deficiency in their immune cells were highly susceptible to *Ct*-CR infection compared with littermate controls **(Figure 2E)**. Vav-Cre is also expressed by endothelial cells (Joseph et al., 2013), which express high levels of AHR (Diny et al., 2022). Therefore, AHRKO^Immune^ mice could simultaneously delete AHR in both their immune and endothelial cells. To account for a potential role for AHR in anti-Cryptosporidial defence via the endothelium only, we made use of a tamoxifen-inducible Cre line controlled by the cadherin 5 (Cdh5) promoter to delete AHR selectively in endothelial cells. Cdh5^CreERT2^AHR^fl/fl^ (AHRKO^Endo^) and littermate control mice were administered tamoxifen orally. Five days post-tamoxifen treatment, (AHRKO^Endo^) mice were infected with *Ct*-CR, alongside a cohort of AHRKO^Immune^ mice in the same experiment. In endothelial cell-specific AHR knockout mice, *C. tyzzeri* levels were comparable to those in littermate controls, whereas the infection burden was again increased in AHRKO^Immune^ mice **(Figure 2F)**. *C. tyzzeri* levels in the ileum of AHRKO^Immune^ mice were also similar to the levels in total body-AHR deficient mice (AHRKO^Body^) (**Figure 2G**). Hence, these results narrowed down the importance of AHR expression specifically in immune cells to control an intestinal *C. tyzzeri* infection. We also noted that the increased parasite burden in AHRKO^Immune^ mice was associated with increased expression of enterocyte markers at the expense of stem cell and Tuft cell markers **(Figure 2H)**. Taken together, AHR signalling in immune cells is vital to regulate the ability of *Cryptosporidium* to grow in intestinal epithelial cells of the small intestine.

**Figure 2:**
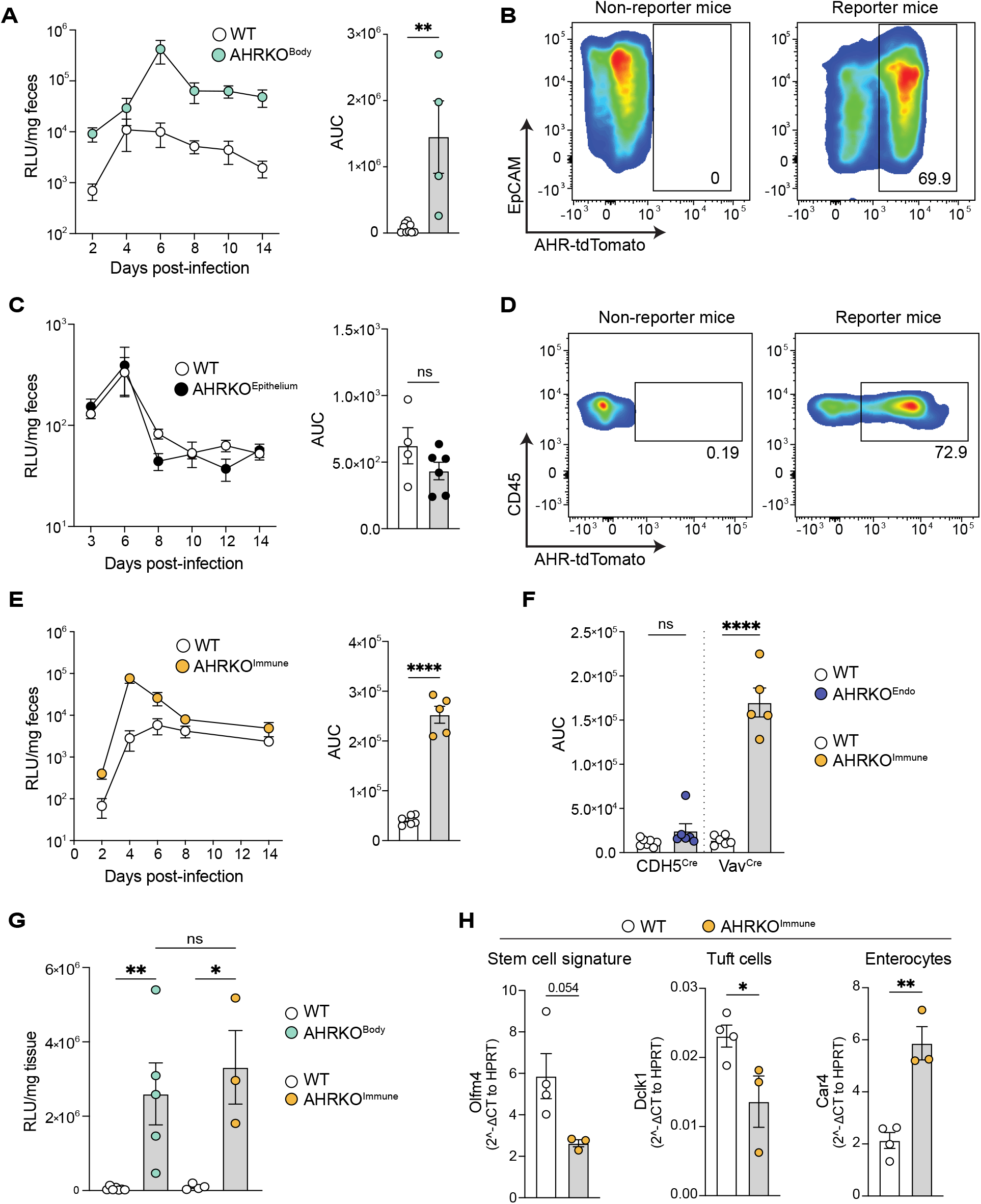
Hematopoietic cell-specific AHR signalling is indispensable to control *C. tyzzeri* infection. (A) *Ct-*CR parasite burden (left panel) and the area under the curve (AUC) (right panel) in WT and total body AHR knockout mice (AHRKO^Body^). (B) AHR-tdTomato expression in EpCAM+ intestinal epithelial cells (C) *Ct-*CR parasite growth in WT (Villin^Cre-^ AHR^fl/fl^) and epithelium-specific AHR knockout mice (Villin^Cre+^ AHR^fl/fl^ termed AHRKO^Epithelium^). (D) AHR- tdTomato expression in small intestinal immune cells (CD45+) (E) *Ct-*CR parasite growth in WT (Vav^Cre-^ AHR^fl/fl^) and immune cell-specific AHR knockout mice (Vav^Cre+^ AHR^fl/fl^ termed AHRKO^Immune^). (F) Area under the curve of *Ct*-CR growth in endothelial (Cdh5^Cre+^ AHR^fl/fl^ termed AHRKO^Endo^) and immune cell-specific AHR knockout mice (AHRKO^Immune^). (G) *Ct-*CR burden at the peak of infection (DPI-6) in total-body (AHRKO^Body^) and immune cell-specific AHR knockout mice (AHRKO^Immune^). (H) qPCR of marker genes of stem cells (*Olfm4*), Tuft cells (*Dclk1*), and enterocytes (*Car4*). (A-H) Representative results of at least 2 independent experiments. Each dot represents individual mice. RLU-Relative Luciferase Units. Error bars, mean + SEM. ns-not significant, *p < 0.05, **p < 0.01, ****p < 0.0001 as calculated by t-test or one-way ANOVA with Tukey post-test.

### AHR-expressing CD8a intraepithelial lymphocytes respond to *C. tyzzeri* parasite infection

Lymphocytes in the small intestine are spatially organised into intraepithelial and lamina propria layers. The intraepithelial layer is enriched with cytotoxic CD8^+^ intraepithelial lymphocytes (IELs), which are the primary responders to epithelial damage due to their close proximity to epithelial cells (Cheroutre et al., 2011). Immunofluorescence images of mouse ileal sections show CD3^+^ IELs in close contact with *C. tyzzeri-*infected epithelial cells **(Figure 3A)**. Using AHR-tdTomato reporter mice, we found that all the major IEL subsets, including TCRαβ^+^CD8αα^+^, TCRαβ^+^CD8αβ^+^, and TCRγδ^+^CD8αα^+^ express AHR-tdTomato **(Figures 3B and Figure S1).** TCRαβ and TCRγδ IELs from *Ct*-CR infected mice had increased expression of Ki67, indicative of a hyperproliferative state **(Figure 3C)**, which corresponded with increased numbers of IELs in *Ct-*CR infected mice **(Figure 3D)**. Moreover, IFNγ expressing CD8^+^ IELs were significantly increased following *Ct-*CR infection (**Figure 3E**). CD8^+^ IELs are quiescent at steady state with minimal proliferation (**Figures 3C**), however they rapidly mount a cytotoxic response to kill target cells through effector proteins such as granzymes (Konjar et al., 2018). Indeed, all 3 major types of CD8^+^ IELs express granzyme-B at steady state (**Figure 3F**). Therefore, IELs sense epithelial invasion by *C. tyzzeri* in a similar manner to other infection settings (Hoytema van Konijnenburg et al., 2017; Konjar et al., 2018) and respond by producing IFNγ and granzyme-B, which are effector mediators of IEL cytotoxicity.

**Figure 3:**
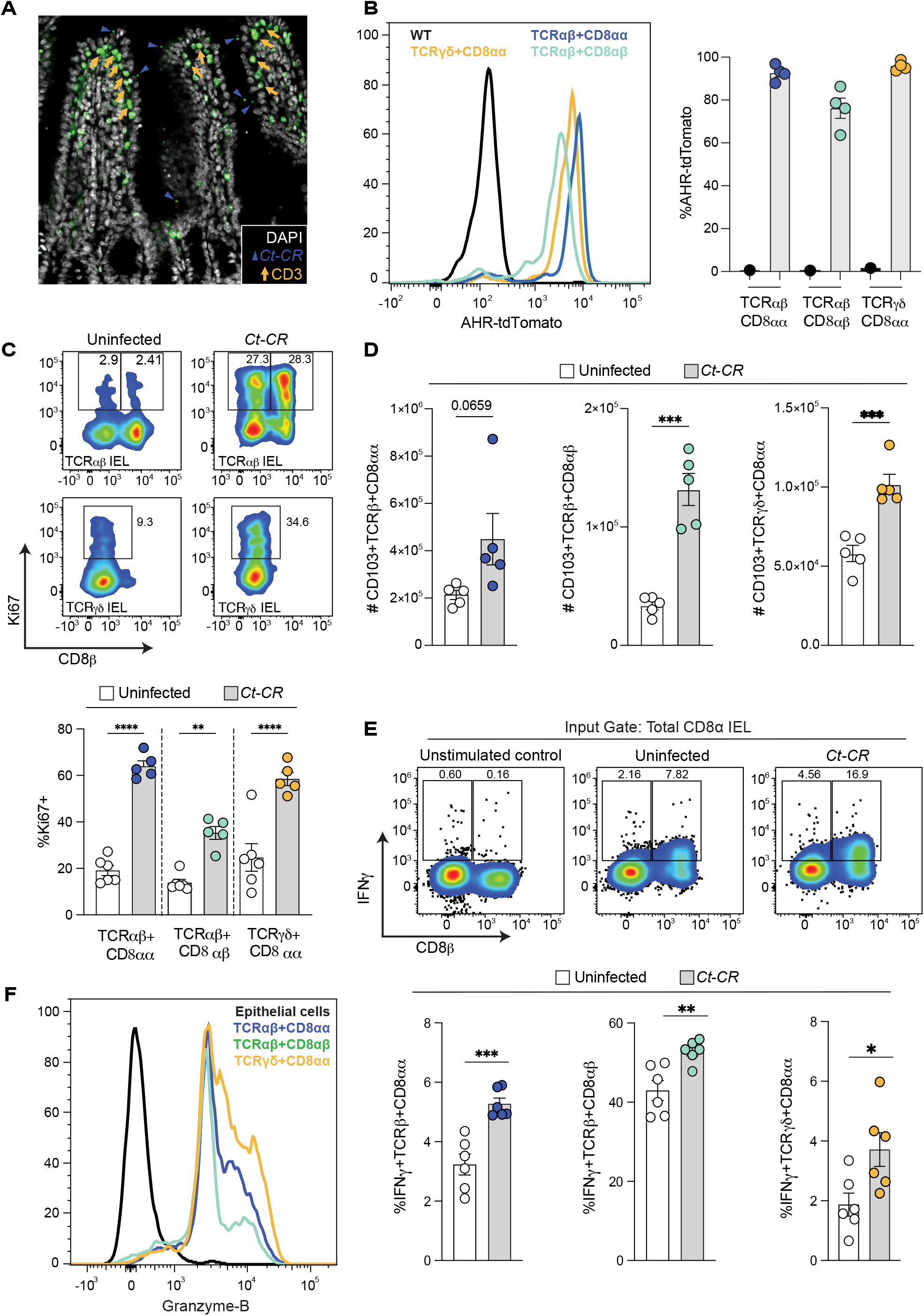
AHR-expressing CD8a intraepithelial lymphocytes respond to *C. tyzzeri* parasite infection. (A) Representative image showing *Ct-*CR (navy blue arrow-head pointing to mNeonGreen positive parasites on the luminal side of villi) and the close proximity of intraepithelial lymphocytes (orange arrow pointing to CD3^+^ IELs) to infected epithelial cells in the ileum. (B) AHR-tdTomato expression by TCRαβ and TCRγδ CD8^+^ IEL subsets. Representative histogram (left) and percentage AHR-tdTomato positive IEL subsets (right) (C) Ki67+ proliferating IELs in *Ct-*CR infected WT mice. Representative FACS plots (top) and percentage Ki67 positive IEL subsets (bottom) (D) Number of IELs in the small intestine of *Ct-* CR infected WT mice. (E) Intracellular IFNγ levels of TCRαβ and TCRγδ CD8^+^ IEL subsets. Representative FACS plots showing IFNγ expressing CD8αβ+ and CD8αα+ IELs (top) and percentage IFNγ+ TCRαβ and TCRγδ CD8^+^ IEL subsets (bottom) in naïve and infected mice. (F) Histogram of intracellular Granzyme-B in TCRαβ and TCRγδ CD8^+^ IEL subsets. CD45 negative epithelial cells are used as negative controls (A-F) Representative results of at least 2 independent experiments. Each dot represents individual mice. Error bars, mean + SEM. *p < 0.05, **p < 0.01, ***p < 0.001, ****p < 0.0001 as calculated by t-test or one-way ANOVA with Tukey post-test.

### AHR-dependent CD8α IELs confer resistance to *C. tyzzeri* infection in immunodeficient mice

AHR expression and signalling is essential for both TCRαβ^+^ and TCRγδ^+^ IEL survival, maintenance, and effector function (Dean et al., 2023; Li et al., 2011; Panda et al., 2023). In line with these findings, the percentage, and total numbers of natural CD8αα IELs recovered from naïve AHRKO^immune^ mice were significantly lower compared to WT littermates whereas the numbers of TCRαβ^+^CD8αβ^+^ IELs were similar **(Figures 4A-4D)**. During infection with *Ct-* CR, TCRαβ^+^CD8αα^+^ and TCRγδ^+^CD8αα^+^ IELs were similarly reduced but TCRαβ^+^CD8αβ^+^ IELs increased in AHRKO^immune^ mice. **(Figures 4E and 4F)**. AHR-deficient CD8α IELs were hyper- proliferative **(Figures 4G and 4H)**. However, they exhibited diminished cytolytic activity indicated by decreased Granzyme-B levels **(Figures 4I and 4J)**. Therefore, it is likely that a decrease in cytotoxic CD8^+^ IELs in AHR deficient mice contributed to the increased *Ct*-CR burden.

**Figure 4:**
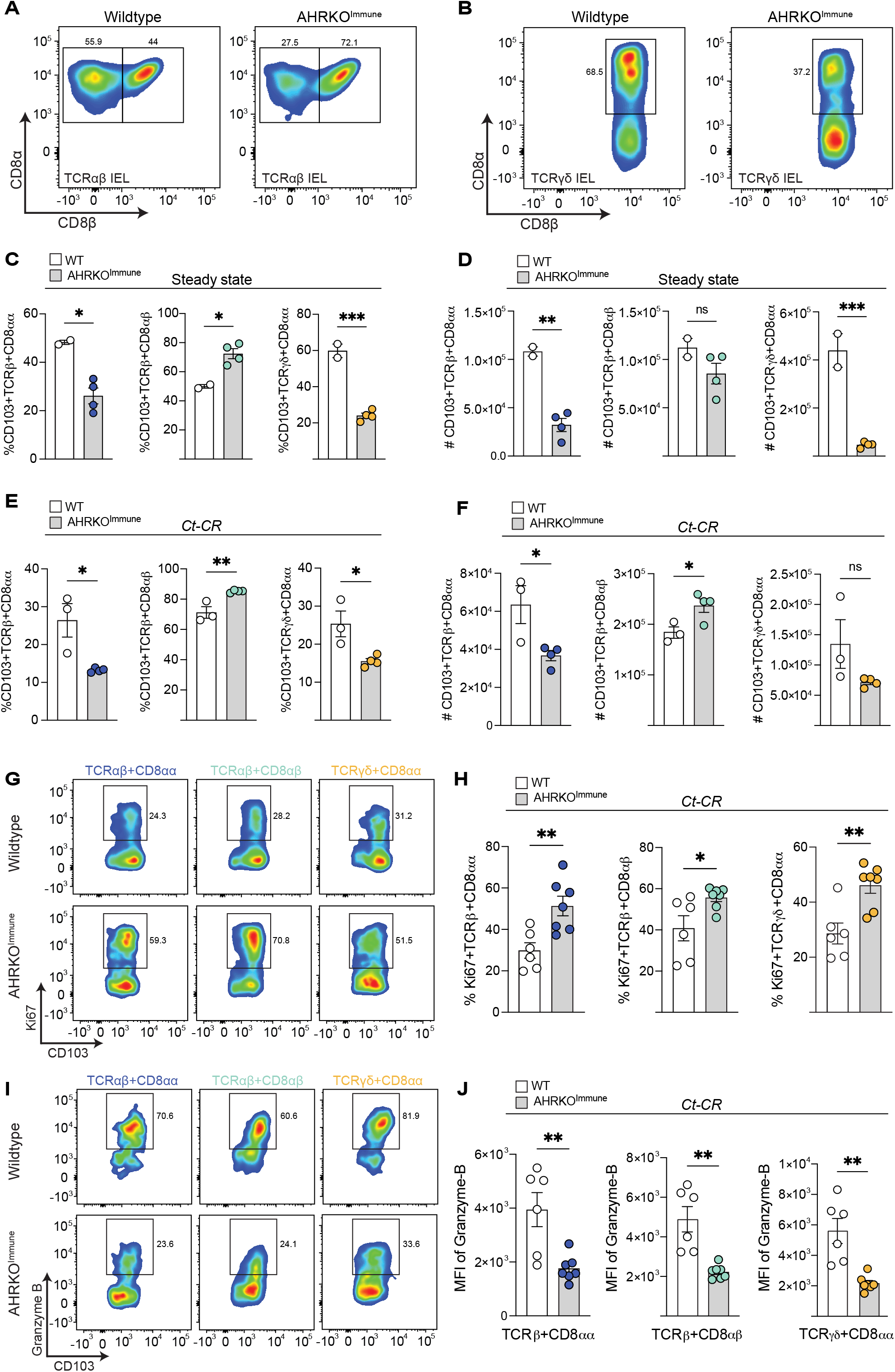
AHR expression influences the CD8a IEL response to *C. tyzzeri* infection. (A & B) Representative FACS plots showing the percentage of TCRαβ^+^CD8αα, TCRαβ^+^CD8αβ IEL subsets (A) and TCRγδ^+^CD8αα (B) in WT and AHRKO^Immune^ mice at steady state. (C & D) Percentage (C) and absolute number (D) of TCRαβ and TCRγδ CD8^+^ IEL subsets in AHRKO^Immune^ mice and littermate WT controls at steady state. (E & F) Percentage (E) and absolute number (F) of CD8^+^ IEL subsets on day 7 post-*Ct-*CR infection. (G & H) Representative FACS plots (G) and percentage (H) of Ki67 positive CD8α IEL subsets in *Ct-*CR-infected mice. (I) Percentage of Granzyme-B-expressing CD8α IEL subsets (J) Mean fluorescence intensity (MFI) of intracellular Granzyme-B in TCRαβ and TCRγδ CD8^+^ IEL subsets in AHRKO^Immune^ mice and littermate WT controls infected with *Ct-*CR. (A-J) Representative results of at least 2 independent experiments. Each dot represents individual mice. Error bars, mean + SEM. ns- not significant, *p < 0.05, **p < 0.01, ***p < 0.001, as calculated by t-test.

To directly assess the protective function of CD8^+^ IELs, we purified CD8^+^ IELs from the small intestine of naïve WT mice and transferred them intravenously into immunodeficient Rag2-IL2Rγ-CD47 triple knockout mice which are usually highly susceptible to infections (Gullicksrud et al., 2022). Four weeks after IEL transfer, the mice were infected with *Ct*-CR **(Figure 5A)**. While untreated immunodeficient mice experienced increased severity with significant reduction in survival **(Figure 5B)**, triple knockout mice that received WT CD8^+^ IELs survived and had reduced parasite burdens in the feces **(Figure 5C)** and ileum **(Figure 5C right panel).** In conclusion, AHR expression in CD8^+^ IELs is required for their maintenance and cytotoxicity, and these CD8^+^ IELs alone are sufficient to control *Ct-*CR parasite growth *in vivo*.

**Figure 5:**
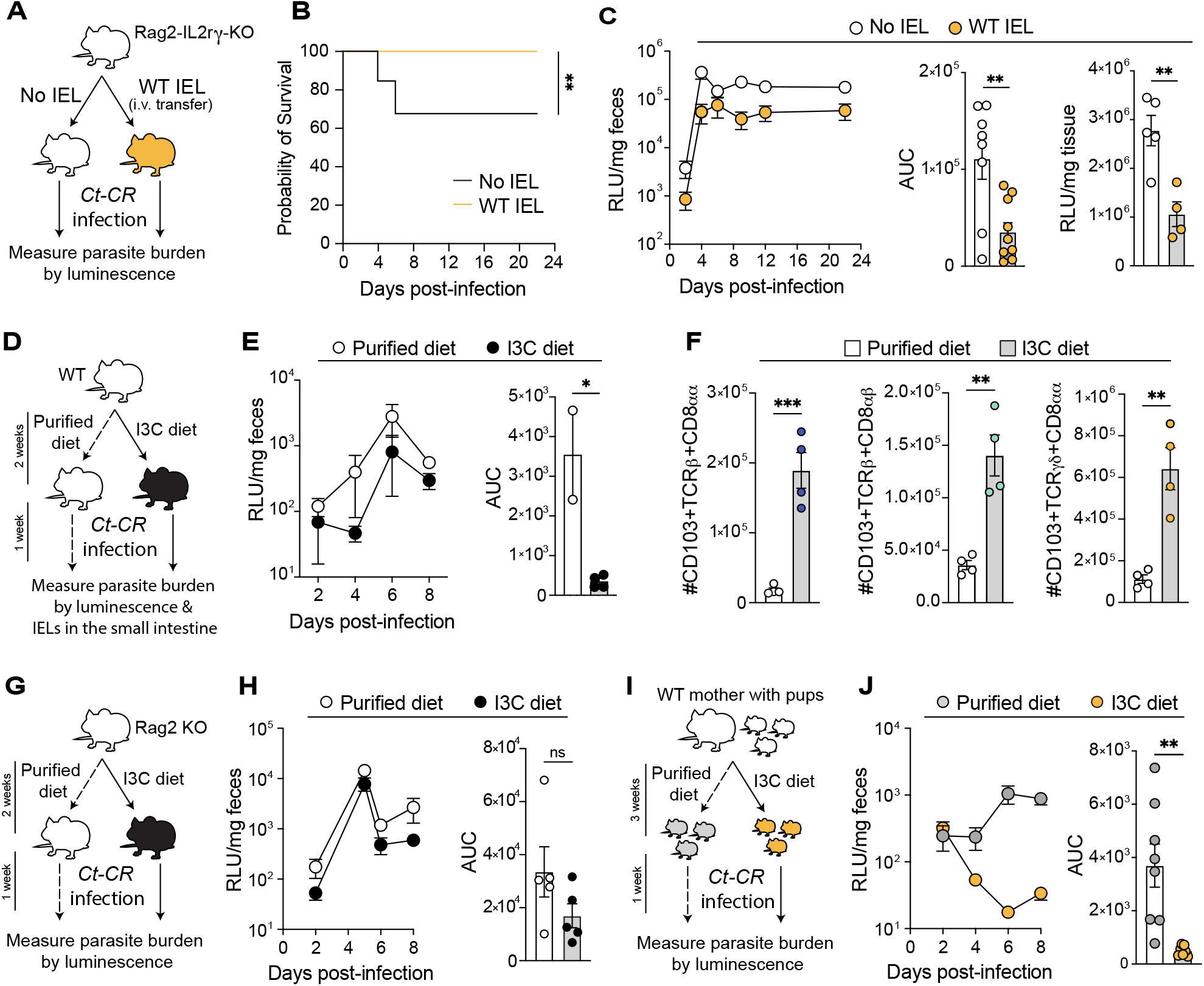
Dietary AHR ligands increase CD8a IELs which confer resistance to *C. tyzzeri* infection. (A) Experimental design of Rag2-IL2Rγ-CD47 triple knockout mice receiving CD8^+^ IELs followed by infection with *Ct-*CR. (B) Survival of Rag2-IL2Rγ-CD47 triple knockout mice infected with *Ct-*CR that received WT CD8^+^ IELs compared to controls. (C) *Ct-*CR parasite burden in the fecal samples of CD8^+^ IEL-transferred Rag2-IL2Rγ-CD47 triple knockout mice. The area under the curve (AUC) is pooled data from 2 independent experiments. *Ct-*CR levels in the ileum on DPI-22 (right panel). (D-F) 3-4 week-old WT mice were fed with either purified diet or I3C diet for 2 weeks followed by infection with *Ct-*CR and maintained on the same diets (D). *Ct-*CR levels in the feces (E) and absolute number of small intestinal CD8α IELs (F) were enumerated. (G & H) Rag2 knockout mice were fed with either purified or I3C diet for 2 weeks and infected with *Ct-*CR while maintained on the same diets (G). Nanoluciferase readings indicating *Ct-*CR burden (H) monitored in the feces (left panel) and AUC (right panel). (I) Experimental design of nursing WT mothers fed with either purified or I3C diets. Their pups were weaned on the same diets and infected with *Ct-*CR. (J) *Ct*-CR parasite burden in young pups of dams that received either purified or I3C diets. (A-F, I & J) Representative results of at least 2 independent experiments. (G & H) Data from 1 experiment. (C) Right panel showing *Ct-*CR levels in the ileum is from 1 experiment. Each dot represents individual mice. Error bars, mean + SEM. ns-not significant, *p < 0.05, **p < 0.01, as calculated by t-test.

### Dietary AHR ligands confer resistance to *C. tyzzeri* infection

AHR is a ligand-activated transcription factor. In the gut, a major source of these ligands are dietary tryptophan-derived phytochemicals and tryptophan metabolites produced by the microbiota(Gomez de Aguero et al., 2016). Since AHR expressing CD8^+^ IELs are key to anti-Cryptosporidial activity, we wanted to determine if this function is influenced by AHR-ligand availability. The phytochemical indole-3-carbinol (I3C), enriched in cruciferous vegetables, is an AHR pro-ligand that is converted to high-affinity AHR ligands upon exposure to stomach acid (Bjeldanes et al., 1991). We first asked whether dietary supplementation with I3C influences CD8^+^ IELs. 3-week-old WT mice were fed either control phytochemical-free synthetic AIN93M diet (‘purified control diet’) or I3C (1000 mg/kg) enriched diet (‘I3C diet’) for 2 weeks and infected with *Ct-*CR while they continued on the same diets (**Figure 5D).** I3C diet fed mice had reduced parasite burdens compared with the mice fed the control diet **(Figure 5E).** Enumeration of CD8^+^ IELs in these mice indicated that I3C diet supplementation robustly increased all subsets of CD8^+^ IELs in the small intestine **(Figure 5F)**. To demonstrate that I3C diet mediated protection from *Ct-*CR infection is T-cell dependent, we fed Rag2 knockout mice with purified control or I3C diet for 2 weeks. Infection with *Ct-*CR resulted in similar burden of parasite in both groups of mice independent of exposure to I3C diet **(Figures 5G and 5H)**. We then investigated the significance of an I3C-enhanced diet in ameliorating *C. tyzzeri* infection in a mouse model that recapitulates early childhood infection in humans. Since AHR signalling is known to impact fertility (Hernandez-Ochoa et al., 2009), pregnant WT females were maintained on normal chow diet until they gave birth. We then changed the diet of WT dams from normal chow to either a purified control diet or an I3C-enriched diet. 3 weeks later, pups were weaned on the same diet that their mothers had received and were infected with *Ct-*CR (**Figure 5I**). WT pups that grew up on the purified control diet were far more susceptible to infection than WT pups that grew up on the I3C diet (**Figure 5J**). Thus, prior exposure to I3C protects mice from *Cryptosporidium* infection.

Taken together, these data underscore the potential of dietary AHR ligands to modulate cytotoxic immune defence against *C. tyzzeri*, leading to improved elimination of infected epithelial cells, and thus offering a therapeutic avenue to prevent or treat cryptosporidiosis by enhancing immunity to *Cryptosporidium* through dietary interventions.

## DISCUSSION

While ubiquitously prevalent, *Cryptosporidium* is particularly problematic in resource-poor settings. Malnutrition is endemic to those same settings, which is in turn a risk factor for *Cryptosporidium* infections and chronic diarrhoea (Iannotti et al., 2015). Malnutrition and immunodeficiency increase the risk of recurrent *Cryptosporidium* infections, which contributes to increased morbidity and mortality (Checkley et al., 2015). The findings presented in this study using a mouse model of infection offers a proof of concept for the potential use of dietary AHR ligands such as I3C to curb this vicious cycle of chronic infections, diarrhoea, and malnutrition in humans and farm animals. Many resource-poor regions across the world already rely on ready-made food formulations such as RUTF (Ready-to-Use Therapeutic Food) to treat children suffering from severe wastage and malnutrition, often as the result of infections, with varying degrees of success (Schoonees et al., 2019). Several human clinical trials show that I3C is safe for human consumption in adults (Naik et al., 2006; Reed et al., 2005; Wong et al., 1997). I3C supplements are commercially available on the market and thus it is conceivable for them to be included in RUTF formulations following controlled human challenge trials confirming their therapeutic potential. Intriguingly, it has also been shown that nursing mice can pass on AHR ligands to their newborn (Lu et al., 2021). Therefore, there is also scope for dietary prophylactic interventions to be made at the level of nursing mothers, especially since *Cryptosporidium* infections are most severe in children less than one year old.

In laboratory settings, younger mice are also more susceptible to *Cryptosporidium* infections. Here, we have shown the importance of cytotoxic CD8^+^ IELs in controlling this infection in an AHR-dependent manner. Due to their close association with intestinal epithelial cells, IELs are known to be primary responders to several invading pathogens in the gut (Edelblum et al., 2015; Hoytema van Konijnenburg et al., 2017; Ismail et al., 2011) and our findings support these observations in the context of *Cryptosporidium* infection. CD8^+^ IELs (natural CD8αα and induced CD8αβ) in the gut are long-lived tissue-resident memory cells. Natural CD8αα cells populate the relatively abiotic guts of pups in response to self-antigens during weaning (Ruscher et al., 2020), while CD8αβ IELs accumulate over time in response to externally-derived antigens (Cheroutre et al., 2011). While the exact mechanism by which these IELs sense epithelial injury during infection is unclear (Hoytema van Konijnenburg et al., 2017), being in a constant state of heightened activation with increased cytolytic granzyme expression (Konjar et al., 2018) increases their ability to swiftly act on *Cryptosporidium*- infected epithelial cells. The presence of IELs at the coalface of the mucosal barrier and their dependence on AHR (Dean et al., 2023; Li et al., 2011; Panda et al., 2023), opens up the possibility to modulate these IELs during *Cryptosporidium* infections through supplementation of dietary AHR ligands.

IFNγ is a key cytokine responsible for early protection from *C. tyzzeri* infection (Gullicksrud et al., 2022; Sateriale et al., 2019). Interestingly, natural CD8αα IELs do not produce as much IFNγ as their induced CD8αβ counterparts although both robustly respond to *C. tyzzeri* infection. However, both TCRαβ^+^ and TCRγδ^+^ IELs expressed cytotoxic granular protein granzyme-B in their quiescent and activated states. A heightened state of activation with increased cytotoxic potential and rapid proliferative capacity could facilitate swift killing of *C. tyzzeri-*infected epithelial cells by IELs. Indeed, AHR deficient mice (AHRKO^Body^ and AHRKO^Immune^) that lack these IELs have elevated parasite burdens from DPI-2 onwards, suggesting that in the absence of cytotoxic CD8^+^ IELs *C. tyzzeri* robustly establish an infection in the small intestine. In contrast to CD8αα IELs, induced CD8αβ IELs can form antigen-specific long-lived memory cells (Soerens et al., 2023). Future studies should explore the antigen-specificity of conventional TCRαβ^+^CD8αβ IELs during primary and secondary infections and narrow down their role in long-term protection against subsequent *Cryptosporidium* infections.

We have previously established *C. tyzzeri* as a model of *Cryptosporidium* infection to study the relationships between a pathogen and its natural host (Sateriale et al., 2019). Our work showed parasite infections caused characteristic villus blunting, a decrease in villus-crypt height ratios, as well as an increase in mitotic events in the epithelium of the small intestine (Sateriale et al., 2019). Here, we further characterised the pathological effects of the *C. tyzzeri*-CR strain on the mouse small intestine and found that it produces many hallmarks of epithelial damage. There is increased expression of *Ifng* at the tissue level, a loss of markers of differentiated cells such as *Chga* and *Dclk1*, as well as an increase in the enterocyte marker *Car4*. Most notably, Sca-1 protein levels significantly increased in infected epithelial tissue, a notable marker of epithelial damage previously seen in mouse models of colitis (Shah et al., 2022; Yui et al., 2018). Increased Sca-1 expression is also caused during infection by the intestinal parasitic helminth *H. polygyrus* in an IFNγ-dependent manner and is indicative of a reversion to a fetal-like state in tissue (Nusse et al., 2018) . We hope that this study paves the way for future work understanding how this parasite specifically damages the small intestine, potentially triggering a re-wiring of the epithelial regenerative response.

Taken together, we have further uncovered host responses to *C. tyzzeri* and revealed the role of environmental sensor AHR and its natural ligands in conferring protection from *Cryptosporidium* infection by modulating gut-resident cytotoxic lymphocytes using a well-defined genetically tractable mouse model. This study extends our understanding of AHR signalling and its importance in maintaining gut barrier integrity in the small intestine and suggests a way forward towards future therapeutic strategies to control the severity of *Cryptosporidium* infections.

## METHODS

### Mouse model of infection

Mice used in this study including C57BL/6, AHR deficient (AHRKO^Body^), AHR^fl/fl^, Villin-Cre, Vav-Cre, CDH5-CreERT2, Rag2 knockout, and Rag2-IL-2rγ-CD47 triple knockout mice were bred in the Francis Crick Institute Biological Research Facility under specific pathogen free conditions. Experiments were conducted as per the guidelines detailed in the Project Licence granted by the UK Home Office. Both male and female mice aged between 3-6 weeks old were randomly assigned to experimental groups. For infection experiments, 50,000 *Ct-*CR oocysts were administered to each mouse by oral gavage. Parasite burden was measured in the feces and intestinal tissues.

### Genome sequencing and alignment

Genomic DNA was isolated from 2x10^7^ *C. tyzzeri* sporozoites using a Qiagen DNeasy Blood & Tissue Kit. A DNA library was prepared with a Nextera XT DNA Library Preparation Kit (Illumina) and 150bp paired end reads were obtained on a MiSeq patform (Illumina). To align the reads, the refence genome for C. tyzzeri (UGA55) was first downloaded from CryptoDB (Amos et al., 2022). Reads were aligned using Burrows-Wheeler Aligner (bwa mem) and converted into bam format using Samtools (Li and Durbin, 2009). Duplicate reads were then removed and reindexed using GATK. Variants were called with GATK HaplotypeCaller using a ploidy of 1 (*Cryptosporidium* sporozoites are haploid) (McKenna et al., 2010). Variants were finally annotated with SnpEff (Cingolani et al., 2012) and plotted against the reference genome using the circlize package for R (Gu et al., 2014).

### Generation of transgenic parasites

Transgenic *C. tyzzeri*-CR parasites were created using methods described previously (Pawlowic et al., 2017; Sateriale et al., 2020). Briefly, 1.5 x 10^6^ oocysts were bleach-treated and incubated at 37 °C in 0.75 % sodium taurocholate for one hour to promote excystation. Sporozoites were transfected with a plasmid expressing Cas9 and guide targeting the *C. tyzzeri* TK gene, along with a repair template carrying Nanoluciferase, mNeonGreen, and Neomycin resistance genes. IFNγ^-/-^ mice were orally gavaged with these transfected sporozoites, following a sodium bicarbonate oral gavage a few minutes prior. Mice were given Paromomycin in their drinking water to select for transgenic parasites and Nanoluciferase levels in fecal samples were tracked over time.

### Measuring parasite burden in tissues and feces

Nanoluciferase readings from fecal samples were performed as a proxy for parasite burdens as previously described using the NanoGlo Luciferase reporter assay (NanoGlo Luciferase kit, Promega, N1150) (Vinayak et al., 2015). Small intestinal tissue sections were cut, weighed, and processed in a similar manner to fecal samples.

### Isolation of parasites

Parasites were isolated from daily mouse fecal collections by sucrose flotation followed by cesium chloride-mediated density gradient centrifugation as described previously (Pawlowic et al., 2017). Briefly, 5-7 days’ worth of fecal samples were made into a slurry in tap water and filtered through a sieve of 250 um pore size. Filtrates were mixed 1:1 with a sucrose solution (1.33 specific gravity) and centrifuged at 1000 g at 4 °C. The supernatant was washed in cold water (1:100), centrifuged again at 1500 g at 4 °C and resuspended in cold saline. This suspension was carefully laid over a cesium chloride solution (1.25 M) in a 1:1 ratio and centrifuged at 16,000 g at 4 °C to produce an interphase containing suspended oocysts. These were collected and stored in cold saline for future use.

### Isolation and transfer of intraepithelial lymphocytes

IELs were isolated from the small intestine. Small intestine was cut open longitudinally and washed twice in Ca^2+^ and Mg^2+^-free PBS to remove the intestinal contents. Cleaned intestine was cut into 1cm pieces and resuspended with HBSS (Ca^2+^ and Mg^2+^-free) containing 5% FCS and 2mM EDTA. Tissue pieces were incubated for 30 minutes at 37°C with shaking at 200 r.p.m. Single cell suspension from the epithelial wash was resuspended in 36% Percoll (Amersham) and layered on top of 67% Percoll (Amersham; Cat# 17-0891-01) and subjected to density gradient centrifugation at room temperature (700g for 30 minutes). Intermediate layer containing IELs was collected and washed with 1X PBS followed by either flowcytometry analysis or used for CD8α IELs purification using EasySep™ mouse CD8α positive selection kit-II (StemCell Technologies; Cat# 18953). Sort purified CD8α IELs (50,000/mouse) were intravenously injected into recipient Rag2-IL-2rγ-CD47 knockout mice.

For measurement of intracellular IFNγ ex-vivo harvested IELs were restimulated with PMA (20ng/ml final concentration) and ionomycin (1μM final concentration) in the presence of brefeldin-A (10μg/ml final concentration) for 3 hours. In some experiments, epithelial cells (top layer of 36% Percoll gradient) that were separated from IELs was used for RNA extraction.

### Flow cytometry

Single cell suspension was first stained with LIVE/DEAD Near-IR dead cell stain (Invitrogen; Cat# L10119). Cell suspensions were then incubated with surface staining antibodies targeting CD45 (Cat# 103146; Clone: 30-F11), CD103 (Cat# 12-1031-82; Clone: 2E7), TCRβ (Cat# 109224; Clone:H57-597), TCRγδ (Cat# 118123 & Cat# 118106; Clone:GL3), CD4 (Cat# 100557; Clone: RM4-5), CD8α (Cat# 100734; Clone:53.6-7), and CD8β (Cat# 740952; Clone: H35-17.2). For intracellular protein staining, cells were fixed and permeabilized using eBioscience FOXP3 staining kit (Invitrogen; Cat# 00-5523-00) followed by intracellular staining of Ki67 (eBiosciences; Cat# 17-5698-82; Clone:SOIA15), Granzyme-B (Biolegend; Cat# 515408; Clone:GB11) and IFNγ (BD Biosciences; Cat# 557724; Clone: XMG1.2) as per the experimental condition. Acquired data was analysed using Flowjo software (https://www.flowjo.com/). Identification of different subsets of IELs was described in the Figure S1.

### Immunohistochemistry and Immunofluorescence Staining

Mouse ileal sections were fixed in 4 % PFA for at least 4 hours and then treated in a solution of 10 % glycerol and 25 % sucrose prior to embedding in OCT. Sections were stained with anti- CD3-AF488 (1: 200; Cat# 100210; Clone: 17A2), EpCAM-AF594 (1:200; Cat# 118222, Clone:G8.8), and DAPI (1:1000; Cat# D9542). Some sections were stained with Sca-1 (1: 200; Cat# 122501; Clone:E13-161.7.) Images were taken with a laser scanning confocal microscope (Zeiss LSM 880) and processed using FIJI (ImageJ version 2.1.0/1.53c).

### Bioimage analysis of Sca-1 immunofluorescence staining

OCT embedded cryosections were stained with Sca-1 as mentioned above. Quantification of Sca-1+ areas in the small intestinal epithelium was performed blindly using QuPath software (Bankhead et al., 2017). The epithelial layer of the villi in the small intestine was delineated manually using the polygon or brush tool in QuPath. 6 villi per small intestine were analyzed. Using the automated quantification tool, mean fluorescence intensity per area unit was estimated. This was repeated in sections stained without primary antibody as controls. The average fluorescence intensity values from the control sections were subtracted from the fluorescence intensity values obtained from each villi in the original samples. The fluorescence intensity values from 6 villi per small intestine were averaged, to report the mean GFP intensity per mouse.

### Quantitative Real-time Polymerase Chain Reaction

RNA from the gut tissues and epithelial cells was extracted using TRIzol RNA purification kit (Invitrogen; Cat# 9738G). cDNA was synthesized with high-Capacity cDNA Reverse Transcription Kit (Thermofisher; Cat# 4368814) and qRT-PCR was performed using Taqman 2x universal PCR master mix (Thermofisher; Cat# 4318157) with appropriate primer sets. Data was normalized to housekeeping genes HPRT or B2M.

### Dietary intervention

For dietary intervention studies mice were fed with synthetic purified control diet (AIN93M) (ssniff; Cat# E15713-047) or AIN93M diet supplemented with indole-3-carbinol (I3C) (1000mg/kg) (ssniff; Cat# S9477-E724) *ad libitum*.

### Statistical analysis

All statistical analysis was performed with GraphPad Prism software (version 9) (https://www.graphpad.com/). For comparison of 2 groups unpaired Student’s *t-*test was used and for multiple comparison analyses a one-way ANOVA followed by Tukey’s multiple comparison test was performed. P <0.05 was considered as significant. Error bars represent mean with standard error mean (SEM). ns-not significant, *p < 0.05, **p < 0.01, ***p < 0.001, ****p < 0.0001.

## Acknowledgements

We would like to acknowledge the Science Technology Platforms at the Francis Crick Institute. We thank the Biological Research Facility for breeding and maintenance of our mouse strains, the Flow Cytometry Facility, the Experimental Histopathology Facility, and the Advanced Light Microscopy Facility. This work was supported by the Francis Crick Institute which receives its core funding from Cancer Research UK (CC2016, CC2063), the UK Medical Research Council (CC2016, CC2063), and the Wellcome Trust (CC2016, CC2063) and a Wellcome Investigator Award to B.S. (210556/Z/18/Z). For the purpose of Open Access, the author has applied a CC BY public copyright licence to any Author Accepted Manuscript version arising from this submission.’

## Author Contributions

M. M., N. B. M., B. S., and A. S. conceived the project, designed experiments, analysed data, and wrote the manuscript. M. M., N. B. M. conducted experiments, analysed data, and prepared figures. O. E. D., T. M., N. L. D., Y. L., A. L., K. S., and M. T., performed experiments or analysed data. M. K. isolated *Ct-*CR2206 and provided advice on parasite propagation.

---

**Figure S1:**
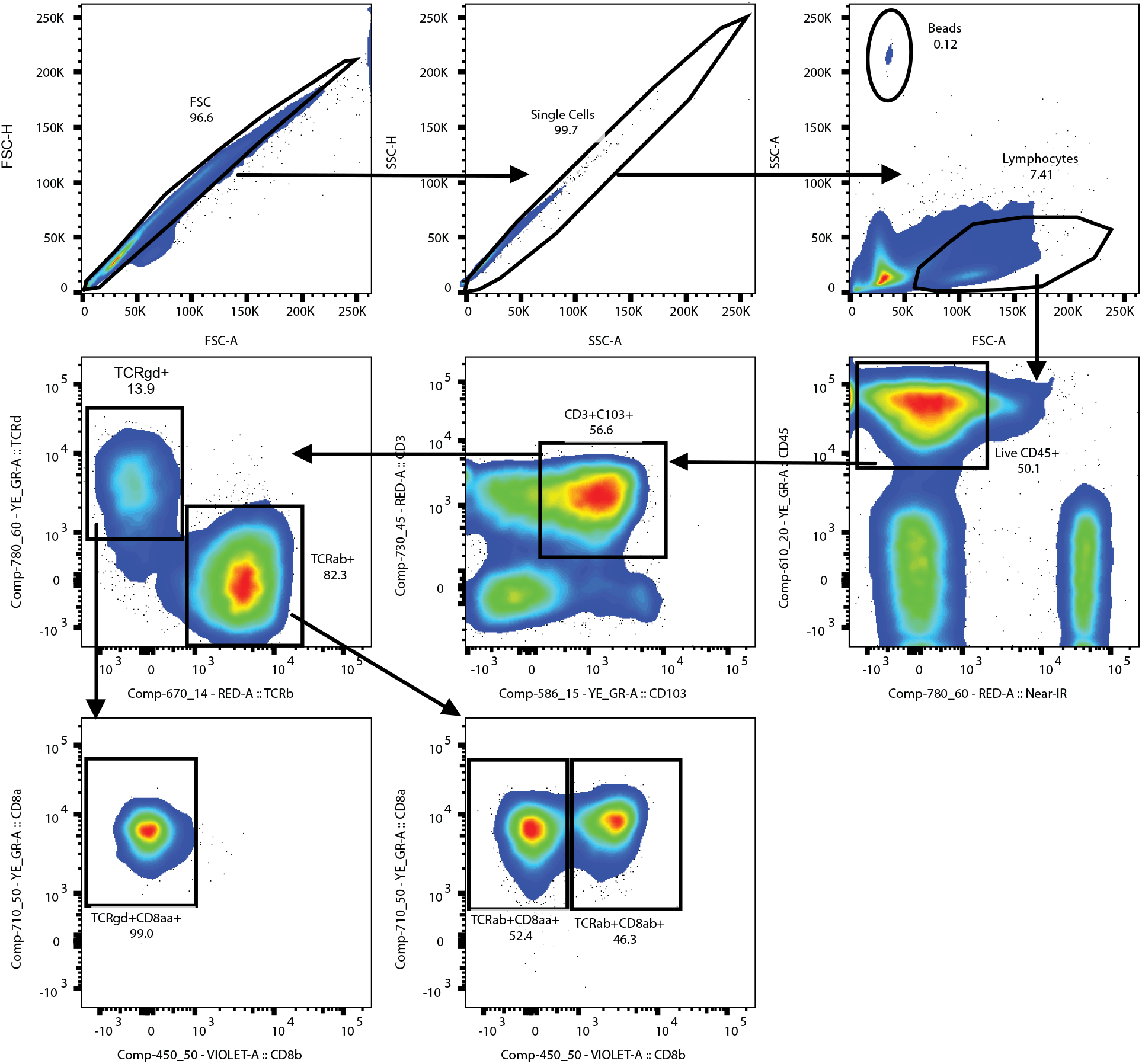
Gating strategy for small intestinal intraepithelial lymphocytes. Single cells were gated based on the forward and side scatter properties. Live CD45^+^ cells that were negative for Live/Dead Near-IR are used to identify CD103^+^ tissue resident lymphocytes. CD103^+^ TCRαβ^+^ CD8αα, CD103^+^ TCRαβ^+^ CD8αβ and CD103^+^ TCRγδ CD8αα IEL subsets were gated from CD45^+^CD103^+^ lymphocytes. CountBright Absolute Counting Beads were used to calculate the absolute number of IELs in the small intestine.

**Table S1:** Summary of genome comparisons between C. tyzzeri strain UGA55 and C. tyzzeri strain CR2206.

|  | SNPs |  | INDELs |
| --- | --- | --- | --- |
| Coding regions | Synonymous | Non-synonymous | 2464 |
|  | 3773 | 5014 |  |
| Non-coding regions | 4315 |  | 518 |

## REFERENCES

Amadi, B., Mwiya, M., Musuku, J., Watuka, A., Sianongo, S., Ayoub, A., and Kelly, P. (2002). Effect of nitazoxanide on morbidity and mortality in Zambian children with cryptosporidiosis: a randomised controlled trial. Lancet 360, 1375–1380.

Amadi, B., Mwiya, M., Sianongo, S., Payne, L., Watuka, A., Katubulushi, M., and Kelly, P. (2009). High dose prolonged treatment with nitazoxanide is not effective for cryptosporidiosis in HIV positive Zambian children: a randomised controlled trial. BMC Infect Dis 9, 195.

Amos, B., Aurrecoechea, C., Barba, M., Barreto, A., Basenko, E.Y., Bazant, W., Belnap, R., Blevins, A.S., Bohme, U., Brestelli, J., et al. (2022). VEuPathDB: the eukaryotic pathogen, vector and host bioinformatics resource center. Nucleic Acids Res 50, D898–D911.

Bankhead, P., Loughrey, M.B., Fernandez, J.A., Dombrowski, Y., McArt, D.G., Dunne, P.D., McQuaid, S., Gray, R.T., Murray, L.J., Coleman, H.G., et al. (2017). QuPath: Open source software for digital pathology image analysis. Sci Rep 7, 16878.

Bjeldanes, L.F., Kim, J.Y., Grose, K.R., Bartholomew, J.C., and Bradfield, C.A. (1991). Aromatic hydrocarbon responsiveness-receptor agonists generated from indole-3-carbinol in vitro and in vivo: comparisons with 2,3,7,8-tetrachlorodibenzo-p-dioxin. Proc Natl Acad Sci U S A 88, 9543–9547.

Chai, J.Y., Guk, S.M., Han, H.K., and Yun, C.K. (1999). Role of intraepithelial lymphocytes in mucosal immune responses of mice experimentally infected with Cryptosporidium parvum. J Parasitol 85, 234–239.

Chappell, C.L., Darkoh, C., Shimmin, L., Farhana, N., Kim, D.K., Okhuysen, P.C., and Hixson, J. (2016). Fecal Indole as a Biomarker of Susceptibility to Cryptosporidium Infection. Infect Immun 84, 2299–2306.

Checkley, W., White, A.C., Jr., Jaganath, D., Arrowood, M.J., Chalmers, R.M., Chen, X.M., Fayer, R., Griffiths, J.K., Guerrant, R.L., Hedstrom, L., et al. (2015). A review of the global burden, novel diagnostics, therapeutics, and vaccine targets for cryptosporidium. Lancet Infect Dis 15, 85–94.

Cheroutre, H., Lambolez, F., and Mucida, D. (2011). The light and dark sides of intestinal intraepithelial lymphocytes. Nat Rev Immunol 11, 445–456.

Cingolani, P., Platts, A., Wang le, L., Coon, M., Nguyen, T., Wang, L., Land, S.J., Lu, X., and Ruden, D.M. (2012). A program for annotating and predicting the effects of single nucleotide polymorphisms, SnpEff: SNPs in the genome of Drosophila melanogaster strain w1118; iso- 2; iso-3. Fly (Austin) 6, 80–92.

Cohn, I.S., Henrickson, S.E., Striepen, B., and Hunter, C.A. (2022). Immunity to Cryptosporidium: Lessons from Acquired and Primary Immunodeficiencies. J Immunol 209, 2261–2268.

Dean, J.W., Helm, E.Y., Fu, Z., Xiong, L., Sun, N., Oliff, K.N., Muehlbauer, M., Avram, D., and Zhou, L. (2023). The aryl hydrocarbon receptor cell intrinsically promotes resident memory CD8(+) T cell differentiation and function. Cell Rep 42, 111963.

Diny, N.L., Schonfeldova, B., Shapiro, M., Winder, M.L., Varsani-Brown, S., and Stockinger, B. (2022). The aryl hydrocarbon receptor contributes to tissue adaptation of intestinal eosinophils in mice. J Exp Med 219.

Edelblum, K.L., Singh, G., Odenwald, M.A., Lingaraju, A., El Bissati, K., McLeod, R., Sperling, A.I., and Turner, J.R. (2015). gammadelta Intraepithelial Lymphocyte Migration Limits Transepithelial Pathogen Invasion and Systemic Disease in Mice. Gastroenterology 148, 1417–1426.

Gomez de Aguero, M., Ganal-Vonarburg, S.C., Fuhrer, T., Rupp, S., Uchimura, Y., Li, H., Steinert, A., Heikenwalder, M., Hapfelmeier, S., Sauer, U., et al. (2016). The maternal microbiota drives early postnatal innate immune development. Science 351, 1296–1302.

Griffiths, J.K., Theodos, C., Paris, M., and Tzipori, S. (1998). The gamma interferon gene knockout mouse: a highly sensitive model for evaluation of therapeutic agents against Cryptosporidium parvum. J Clin Microbiol 36, 2503–2508.

Gu, Z., Gu, L., Eils, R., Schlesner, M., and Brors, B. (2014). circlize Implements and enhances circular visualization in R. Bioinformatics 30, 2811–2812.

Guerin, A., and Striepen, B. (2020). The Biology of the Intestinal Intracellular Parasite Cryptosporidium. Cell Host Microbe 28, 509–515.

Gullicksrud, J.A., Sateriale, A., Engiles, J.B., Gibson, A.R., Shaw, S., Hutchins, Z.A., Martin, L., Christian, D.A., Taylor, G.A., Yamamoto, M., et al. (2022). Enterocyte-innate lymphoid cell crosstalk drives early IFN-gamma-mediated control of Cryptosporidium. Mucosal Immunol 15, 362–372.

Hayday, A., Theodoridis, E., Ramsburg, E., and Shires, J. (2001). Intraepithelial lymphocytes: exploring the Third Way in immunology. Nat Immunol 2, 997–1003.

Heine, J., Moon, H.W., and Woodmansee, D.B. (1984). Persistent Cryptosporidium infection in congenitally athymic (nude) mice. Infect Immun 43, 856–859.

Hernandez-Ochoa, I., Karman, B.N., and Flaws, J.A. (2009). The role of the aryl hydrocarbon receptor in the female reproductive system. Biochem Pharmacol 77, 547–559.

Hossain, M.J., Saha, D., Antonio, M., Nasrin, D., Blackwelder, W.C., Ikumapayi, U.N., Mackenzie, G.A., Adeyemi, M., Jasseh, M., Adegbola, R.A., et al. (2019). Cryptosporidium infection in rural Gambian children: Epidemiology and risk factors. PLoS Negl Trop Dis 13, e0007607.

Hoytema van Konijnenburg, D.P., and Mucida, D. (2017). Intraepithelial lymphocytes. Curr Biol 27, R737–R739.

Hoytema van Konijnenburg, D.P., Reis, B.S., Pedicord, V.A., Farache, J., Victora, G.D., and Mucida, D. (2017). Intestinal Epithelial and Intraepithelial T Cell Crosstalk Mediates a Dynamic Response to Infection. Cell 171, 783–794 e713.

Iannotti, L.L., Trehan, I., Clitheroe, K.L., and Manary, M.J. (2015). Diagnosis and treatment of severely malnourished children with diarrhoea. J Paediatr Child Health 51, 387–395.

Ismail, A.S., Severson, K.M., Vaishnava, S., Behrendt, C.L., Yu, X., Benjamin, J.L., Ruhn, K.A., Hou, B., DeFranco, A.L., Yarovinsky, F., and Hooper, L.V. (2011). Gammadelta intraepithelial lymphocytes are essential mediators of host-microbial homeostasis at the intestinal mucosal surface. Proc Natl Acad Sci U S A 108, 8743–8748.

Joseph, C., Quach, J.M., Walkley, C.R., Lane, S.W., Lo Celso, C., and Purton, L.E. (2013). Deciphering hematopoietic stem cells in their niches: a critical appraisal of genetic models, lineage tracing, and imaging strategies. Cell Stem Cell 13, 520–533.

Kabir, M., Alam, M., Nayak, U., Arju, T., Hossain, B., Tarannum, R., Khatun, A., White, J.A., Ma, J.Z., Haque, R., et al. (2021). Nonsterile immunity to cryptosporidiosis in infants is associated with mucosal IgA against the sporozoite and protection from malnutrition. PLoS Pathog 17, e1009445.

Kaplan, J.E., Benson, C., Holmes, K.K., Brooks, J.T., Pau, A., Masur, H., Centers for Disease, C., Prevention, National Institutes of, H., and America, H.I.V.M.A.o.t.I.D.S.o. (2009). Guidelines for prevention and treatment of opportunistic infections in HIV-infected adults and adolescents: recommendations from CDC, the National Institutes of Health, and the HIV Medicine Association of the Infectious Diseases Society of America. MMWR Recomm Rep 58, 1–207; quiz CE201-204.

Kattula, D., Jeyavelu, N., Prabhakaran, A.D., Premkumar, P.S., Velusamy, V., Venugopal, S., Geetha, J.C., Lazarus, R.P., Das, P., Nithyanandhan, K., et al. (2017). Natural History of Cryptosporidiosis in a Birth Cohort in Southern India. Clin Infect Dis 64, 347–354.

Khalil, I.A., Troeger, C., Rao, P.C., Blacker, B.F., Brown, A., Brewer, T.G., Colombara, D.V., De Hostos, E.L., Engmann, C., Guerrant, R.L., et al. (2018). Morbidity, mortality, and long-term consequences associated with diarrhoea from Cryptosporidium infection in children younger than 5 years: a meta-analyses study. Lancet Glob Health 6, e758–e768.

Konjar, S., Frising, U.C., Ferreira, C., Hinterleitner, R., Mayassi, T., Zhang, Q., Blankenhaus, B., Haberman, N., Loo, Y., Guedes, J., et al. (2018). Mitochondria maintain controlled activation state of epithelial-resident T lymphocytes. Sci Immunol 3.

Kotloff, K.L., Nataro, J.P., Blackwelder, W.C., Nasrin, D., Farag, T.H., Panchalingam, S., Wu, Y., Sow, S.O., Sur, D., Breiman, R.F., et al. (2013). Burden and aetiology of diarrhoeal disease in infants and young children in developing countries (the Global Enteric Multicenter Study, GEMS): a prospective, case-control study. Lancet 382, 209–222.

Kvac, M., McEvoy, J., Loudova, M., Stenger, B., Sak, B., Kvetonova, D., Ditrich, O., Raskova, V., Moriarty, E., Rost, M., et al. (2013). Coevolution of Cryptosporidium tyzzeri and the house mouse (Mus musculus). Int J Parasitol 43, 805–817.

Li, H., and Durbin, R. (2009). Fast and accurate short read alignment with Burrows-Wheeler transform. Bioinformatics 25, 1754–1760.

Li, Y., Innocentin, S., Withers, D.R., Roberts, N.A., Gallagher, A.R., Grigorieva, E.F., Wilhelm, C., and Veldhoen, M. (2011). Exogenous stimuli maintain intraepithelial lymphocytes via aryl hydrocarbon receptor activation. Cell 147, 629–640.

Liu, L., Johnson, H.L., Cousens, S., Perin, J., Scott, S., Lawn, J.E., Rudan, I., Campbell, H., Cibulskis, R., Li, M., et al. (2012). Global, regional, and national causes of child mortality: an updated systematic analysis for 2010 with time trends since 2000. Lancet 379, 2151–2161.

Lu, P., Yamaguchi, Y., Fulton, W.B., Wang, S., Zhou, Q., Jia, H., Kovler, M.L., Salazar, A.G., Sampah, M., Prindle, T., Jr., et al. (2021). Maternal aryl hydrocarbon receptor activation protects newborns against necrotizing enterocolitis. Nat Commun 12, 1042.

Manabe, Y.C., Clark, D.P., Moore, R.D., Lumadue, J.A., Dahlman, H.R., Belitsos, P.C., Chaisson, R.E., and Sears, C.L. (1998). Cryptosporidiosis in patients with AIDS: correlates of disease and survival. Clin Infect Dis 27, 536–542.

Manjunatha, U.H., Vinayak, S., Zambriski, J.A., Chao, A.T., Sy, T., Noble, C.G., Bonamy, G.M.C., Kondreddi, R.R., Zou, B., Gedeck, P., et al. (2017). A Cryptosporidium PI(4)K inhibitor is a drug candidate for cryptosporidiosis. Nature 546, 376–380.

Marzook, N.B., and Sateriale, A. (2020). Crypto-Currency: Investing in New Models to Advance the Study of Cryptosporidium Infection and Immunity. Front Cell Infect Microbiol 10, 587296.

McKenna, A., Hanna, M., Banks, E., Sivachenko, A., Cibulskis, K., Kernytsky, A., Garimella, K., Altshuler, D., Gabriel, S., Daly, M., and DePristo, M.A. (2010). The Genome Analysis Toolkit: a MapReduce framework for analyzing next-generation DNA sequencing data. Genome Res 20, 1297–1303.

Metidji, A., Omenetti, S., Crotta, S., Li, Y., Nye, E., Ross, E., Li, V., Maradana, M.R., Schiering, C., and Stockinger, B. (2018). The Environmental Sensor AHR Protects from Inflammatory Damage by Maintaining Intestinal Stem Cell Homeostasis and Barrier Integrity. Immunity 49, 353–362 e355.

Molbak, K., Andersen, M., Aaby, P., Hojlyng, N., Jakobsen, M., Sodemann, M., and da Silva, A.P. (1997). Cryptosporidium infection in infancy as a cause of malnutrition: a community study from Guinea-Bissau, west Africa. Am J Clin Nutr 65, 149–152.

Naik, R., Nixon, S., Lopes, A., Godfrey, K., Hatem, M.H., and Monaghan, J.M. (2006). A randomized phase II trial of indole-3-carbinol in the treatment of vulvar intraepithelial neoplasia. Int J Gynecol Cancer 16, 786–790.

Nusse, Y.M., Savage, A.K., Marangoni, P., Rosendahl-Huber, A.K.M., Landman, T.A., de Sauvage, F.J., Locksley, R.M., and Klein, O.D. (2018). Parasitic helminths induce fetal-like reversion in the intestinal stem cell niche. Nature 559, 109–113.

Panda, S.K., Peng, V., Sudan, R., Ulezko Antonova, A., Di Luccia, B., Ohara, T.E., Fachi, J.L., Grajales-Reyes, G.E., Jaeger, N., Trsan, T., et al. (2023). Repression of the aryl-hydrocarbon receptor prevents oxidative stress and ferroptosis of intestinal intraepithelial lymphocytes. Immunity.

Pasquali, P., Fayer, R., Almeria, S., Trout, J., Polidori, G.A., and Gasbarre, L.C. (1997). Lymphocyte dynamic patterns in cattle during a primary infection with Cryptosporidium parvum. J Parasitol 83, 247–250.

Pawlowic, M.C., Vinayak, S., Sateriale, A., Brooks, C.F., and Striepen, B. (2017). Generating and Maintaining Transgenic Cryptosporidium parvum Parasites. Curr Protoc Microbiol 46, 20B 22 21-20B 22 32.

Reed, G.A., Peterson, K.S., Smith, H.J., Gray, J.C., Sullivan, D.K., Mayo, M.S., Crowell, J.A., and Hurwitz, A. (2005). A phase I study of indole-3-carbinol in women: tolerability and effects. Cancer Epidemiol Biomarkers Prev 14, 1953–1960.

Robinson, P., Okhuysen, P.C., Chappell, C.L., Lewis, D.E., Shahab, I., Lahoti, S., and White, A.C., Jr. (2001). Expression of IL-15 and IL-4 in IFN-gamma-independent control of experimental human Cryptosporidium parvum infection. Cytokine 15, 39–46.

Ruscher, R., Lee, S.T., Salgado, O.C., Breed, E.R., Osum, S.H., and Hogquist, K.A. (2020). Intestinal CD8alphaalpha IELs derived from two distinct thymic precursors have staggered ontogeny. J Exp Med 217.

Russler-Germain, E.V., Jung, J., Miller, A.T., Young, S., Yi, J., Wehmeier, A., Fox, L.E., Monte, K.J., Chai, J.N., Kulkarni, D.H., et al. (2021). Commensal Cryptosporidium colonization elicits a cDC1-dependent Th1 response that promotes intestinal homeostasis and limits other infections. Immunity 54, 2547–2564 e2547.

Sasson, S.C., Gordon, C.L., Christo, S.N., Klenerman, P., and Mackay, L.K. (2020). Local heroes or villains: tissue-resident memory T cells in human health and disease. Cell Mol Immunol 17, 113–122.

Sateriale, A., Pawlowic, M., Vinayak, S., Brooks, C., and Striepen, B. (2020). Genetic Manipulation of Cryptosporidium parvum with CRISPR/Cas9. Methods Mol Biol 2052, 219–228.

Sateriale, A., Slapeta, J., Baptista, R., Engiles, J.B., Gullicksrud, J.A., Herbert, G.T., Brooks, C.F., Kugler, E.M., Kissinger, J.C., Hunter, C.A., and Striepen, B. (2019). A Genetically Tractable, Natural Mouse Model of Cryptosporidiosis Offers Insights into Host Protective Immunity. Cell Host Microbe 26, 135–146 e135.

Schindelin, J., Arganda-Carreras, I., Frise, E., Kaynig, V., Longair, M., Pietzsch, T., Preibisch, S., Rueden, C., Saalfeld, S., Schmid, B., et al. (2012). Fiji: an open-source platform for biological-image analysis. Nat Methods 9, 676–682.

Schoonees, A., Lombard, M.J., Musekiwa, A., Nel, E., and Volmink, J. (2019). Ready-to-use therapeutic food (RUTF) for home-based nutritional rehabilitation of severe acute malnutrition in children from six months to five years of age. Cochrane Database Syst Rev 5, CD009000.

Sepahvand, F., Mamaghani, A.J., Ezatpour, B., Badparva, E., Zebardast, N., and Fallahi, S. (2022). Gastrointestinal parasites in immunocompromised patients; A comparative cross-sectional study. Acta Trop 231, 106464.

Shah, K., Maradana, M.R., Joaquina Delas, M., Metidji, A., Graelmann, F., Llorian, M., Chakravarty, P., Li, Y., Tolaini, M., Shapiro, M., et al. (2022). Cell-intrinsic Aryl Hydrocarbon Receptor signalling is required for the resolution of injury-induced colonic stem cells. Nat Commun 13, 1827.

Soerens, A.G., Kunzli, M., Quarnstrom, C.F., Scott, M.C., Swanson, L., Locquiao, J.J., Ghoneim, H.E., Zehn, D., Youngblood, B., Vezys, V., and Masopust, D. (2023). Functional T cells are capable of supernumerary cell division and longevity. Nature 614, 762–766.

Stockinger, B., Shah, K., and Wincent, E. (2021). AHR in the intestinal microenvironment: safeguarding barrier function. Nat Rev Gastroenterol Hepatol 18, 559–570.

Sujino, T., London, M., Hoytema van Konijnenburg, D.P., Rendon, T., Buch, T., Silva, H.M., Lafaille, J.J., Reis, B.S., and Mucida, D. (2016). Tissue adaptation of regulatory and intraepithelial CD4(+) T cells controls gut inflammation. Science 352, 1581–1586.

Vinayak, S., Pawlowic, M.C., Sateriale, A., Brooks, C.F., Studstill, C.J., Bar-Peled, Y., Cipriano, M.J., and Striepen, B. (2015). Genetic modification of the diarrhoeal pathogen Cryptosporidium parvum. Nature 523, 477–480.

Wong, G.Y., Bradlow, L., Sepkovic, D., Mehl, S., Mailman, J., and Osborne, M.P. (1997). Dose-ranging study of indole-3-carbinol for breast cancer prevention. J Cell Biochem Suppl 28-29, 111-116.

Yui, S., Azzolin, L., Maimets, M., Pedersen, M.T., Fordham, R.P., Hansen, S.L., Larsen, H.L., Guiu, J., Alves, M.R.P., Rundsten, C.F., et al. (2018). YAP/TAZ-Dependent Reprogramming of Colonic Epithelium Links ECM Remodeling to Tissue Regeneration. Cell Stem Cell 22, 35–49 e37.

